# Comparative analysis of new, mScarlet-based red fluorescent tags in *Caenorhabditis elegans*

**DOI:** 10.1101/2024.06.11.598534

**Authors:** Wen Xi Cao, Daniel Merritt, Karinna Pe, Michael Cesar, Oliver Hobert

**Author notes:** Corresponding authors: (WXC), (OH).

## Abstract

One problem that has hampered the use of red fluorescent proteins in the fast-developing nematode *C. elegans* has been the substantial time delay in maturation of several generations of red fluorophores. The recently described mScarlet-I3 protein has properties that may overcome this limitation. We compare here the brightness and maturation time of CRISPR/Cas9 genome-engineered mScarlet, mScarlet3, mScarlet-I3 and GFP reporter knock-ins. Comparing the onset and brightness of expression of reporter alleles of *C. elegans golg-4*, encoding a broadly expressed Golgi resident protein, we found that the onset of detection of mScarlet-I3 in the embryo is several hours earlier than older versions of mScarlet and comparable to GFP. These findings were further supported by comparing mScarlet-I3 and GFP reporter alleles for *pks-1*, a gene expressed in the CAN neuron and cells of the alimentary system, as well as reporter alleles for the panneuronal, nuclear marker *unc-75*. Hence, the relative properties of mScarlet-I3 and GFP do not depend on cellular or subcellular context. In all cases, mScarlet-I3 reporters also show improved signal-to-noise ratio compared to GFP.

## INTRODUCTION

mScarlet is a commonly used fluorophore that provides many advantages over previous red fluorescent proteins (RFPs) (BINDELS *et al*. 2017). In *C. elegans*, mScarlet (also known as wrmScarlet) has been shown to be eight-fold brighter than TagRFP-T, to show improved signal-to-noise ratio, and to be more red-shifted than TagRFP-T (EL MOURIDI *et al*. 2017). These properties maximize signal while limiting bleed-through when used in conjunction with a green or yellow-green fluorophore such as GFP or mNeonGreen (EL MOURIDI *et al*. 2017). Furthermore, mScarlet is specifically designed to be monomeric, and does not pass through a green immature intermediate like other red fluorophores, such as tdTomato or mCherry (WIEHLER *et al*. 2001; VERKHUSHA *et al*. 2004; WACHTER *et al*. 2010; BINDELS *et al*. 2017).

However, a lingering issue that plagues most RFPs used in *C. elegans*, including mScarlet, is their slow maturation time. Green fluorophores, including GFP, mEGFP and mNeonGreen are both rapidly-maturing and bright, with characterized maturation times of around 30 minutes or less at 37°C in cell culture [(BALLEZA *et al*. 2018); *fpbase*.*org*].

In contrast, many RFPs display maturation times on the order of several hours at 37°C in cell culture [(BALLEZA *et al*. 2018); *fpbase*.*org*]. In the rapidly developing model organism *C. elegans*, where key developmental events occur over time scales of very few hours (e.g. the cell cycle time in the developing embryo is on the order of ∼20 minutes, and complete embryo hatches after ∼800 minutes of development), delayed fluorophore maturation impedes efforts to visualize rapid changes in gene expression. Also, the fluorophore-based visualization of cellular morphology changes often necessitates the availability of rapidly maturing fluorophores. For example, given that many cell type-selective drivers only become active during terminal differentiation, slowly maturing RFPs cannot be used to visualize processes, such as neurite outgrowth, which only happen at or around the time of terminal differentiation.

Recently, rapidly-maturing derivatives of mScarlet have been described (GADELLA *et al*. 2023). These RFPs, namely mScarlet3 and mScarlet-I3, have been characterized in cells lines to have much improved maturation times and completeness with minimal impact on brightness, and to retain nearly identical excitation/emission spectra as mScarlet. When characterized *in vivo* and in mammalian cell culture, mScarlet3 was found to be brighter while maturing four-times faster than mScarlet (GADELLA *et al*. 2023). mScarlet-I3 is the fastest maturing RFP characterized with only a slight decrease in relative brightness when compared to mScarlet (GADELLA *et al*. 2023). Here, we codon-optimize these new RFPs, generate a series of reporter alleles via CRISPR/Cas9-mediated engineering into *C. elegans* genome, and compare their intensity and dynamics *in vivo* against currently-used mScarlet and GFP reporters.

## MATERIALS AND METHODS

### C. elegans strains and maintenance

*C. elegans* strains were cultivated at 20°C on plates containing Nematode Growth Media seeded with *E. coli* OP50.

OH19204 *golg-4(ot1508[gfp::golg-4])*

PHX6547 *golg-4(syb6547[wrmScarlet::golg-4)*

OH19205 *golg-4(ot1509[mScarlet3::golg-4])*

OH19206 *golg-4(ot1510[mScarlet-I3::golg-4])*

OH19124 *pks-1(ot1488[pks-1::SL2::mScarlet-I3::h2b])*

OH19125 *pks-1(ot1489[pks-1::SL2::gfp::h2b])*

PHX6499 *unc-75(syb6499[gfp::unc-75])*

OH19299 *unc-75(ot1539[mScarlet-I3::unc-75])*

### Fluorophore design and transgenics

Sequences of the fluorophore tags used can be found in Supplemental File S1. *C. elegans* codon-optimized GFP coding sequence containing three introns was amplified from pPD95.75 plasmid. Protein sequences of mScarlet3 and mScarlet-I3 were obtained from *fpbase*.*org* and (GADELLA *et al*. 2023), and reverse translated. The coding sequence were codon-optimized for *C. elegans* expression with two introns added, using sequences recommended by the MPI *C. elegans* codon adapter (REDEMANN *et al*. 2011), and synthesized by IDT. Fluorophores were knocked into the respective genomic loci using CRISPR/Cas9-based genome editing protocol described in (EROGLU *et al*. 2023), using *dpy-10* co-conversion marker and reagents from IDT. PHX6547 and PHX6499 were generated by SunyBiotech.

### Embryo collection and staging

Embryos for microscopy imaging were obtained either by directly picking off the plates onto microscope slides with agar pads (for later stages), dissection of gravid adults (for earlier stages), or collected in large numbers with a gentle egg prep. To dissect, gravid adults were transferred into a droplet of M9 on a glass slide, and cut through the midbody using an 18-gauge needle. Released embryos were aspirated into a microcapillary pipette and transferred directly onto agar pad slides by mouth pipette. For the gentle egg prep, worms were washed off nearly starved plates enriched for embryos and gravid adults with M9 buffer and pelleted by centrifugation at 2000 x g for 1 minute. Worms were treated in a mixture of 500μl distilled water, 500μl household bleach, and 50μl of 10M sodium hydroxide for 4-5 minutes with shaking, or just until adult bodies break apart and eggs are released. Embryos are pelleted by centrifugation and washed with 1ml of M9 three times, then resuspended in M9 buffer and spotted onto agar pad slides for imaging. Individual embryo stages were visualized by DIC at the time of imaging.

### Microscopy

Adult worms were mounted on 5% agar pads and anesthetized with 50mM sodium azide in M9 buffer. Embryos were obtained as described above, and mounted in M9 buffer without sodium azide. Images were acquired on a Zeiss LSM980 confocal microscope (*golg-4* and *unc-75* strains) or the Zeiss Axioimager Z2 light microscope (*pks-1* strains) using the ZEN Blue image acquisition software with preset excitation and emission parameters for EGFP and mScarlet (for all red fluorophore variants). For each image, Z-stacks of 1μm thickness were obtained for the entire sample, and the medial section was later chosen for analysis.

### Image quantification and analyses

For embryos, representative images of the approximate medial section of embryos acquired at each stage are shown. For adult expression, fluorescence intensity of a single Z-plane imaged through the middle of the adult grinder, visualized by DIC, were chosen for quantification. For representative images of UNC-75 expression in adults, maximum intensity projections were shown to visualize all neuron nuclei. Images were analyzed using ImageJ FIJI (SCHINDELIN *et al*. 2012) and quantifications were visualized using GraphPad PRISM 8.3.0. For each fluorophore, average fluorescence intensity of the ROI of at least 10 individual animals are quantified plotted, and the median values are compared.

## RESULTS AND DISCUSSION

### mScarlet3 is an exceptionally bright fluorophore in *C. elegans*

To assess the brightness of the new RFP variants, mScarlet3 and mScarlet-I3, against mScarlet (aka wrmScarlet) in *C. elegans*, we designed *C. elegans* codon-optimized, artificial intron-containing mScarlet3 and mScarlet-I3 coding sequences. Using CRISPR/Cas9 genome editing, these fluorophores were inserted into the N-terminus of ubiquitously-expressed GOLG-4, the *C. elegans* ortholog of a trans Golgi-associated coiled-coil golgin protein golgin-245 (MUNRO 2011). Expression of these fluorophores were analyzed in parallel with and compared to mScarlet expression (wrmScarlet::GOLG-4), also inserted into the same location (**Figure 1A**).

**Figure 1.**
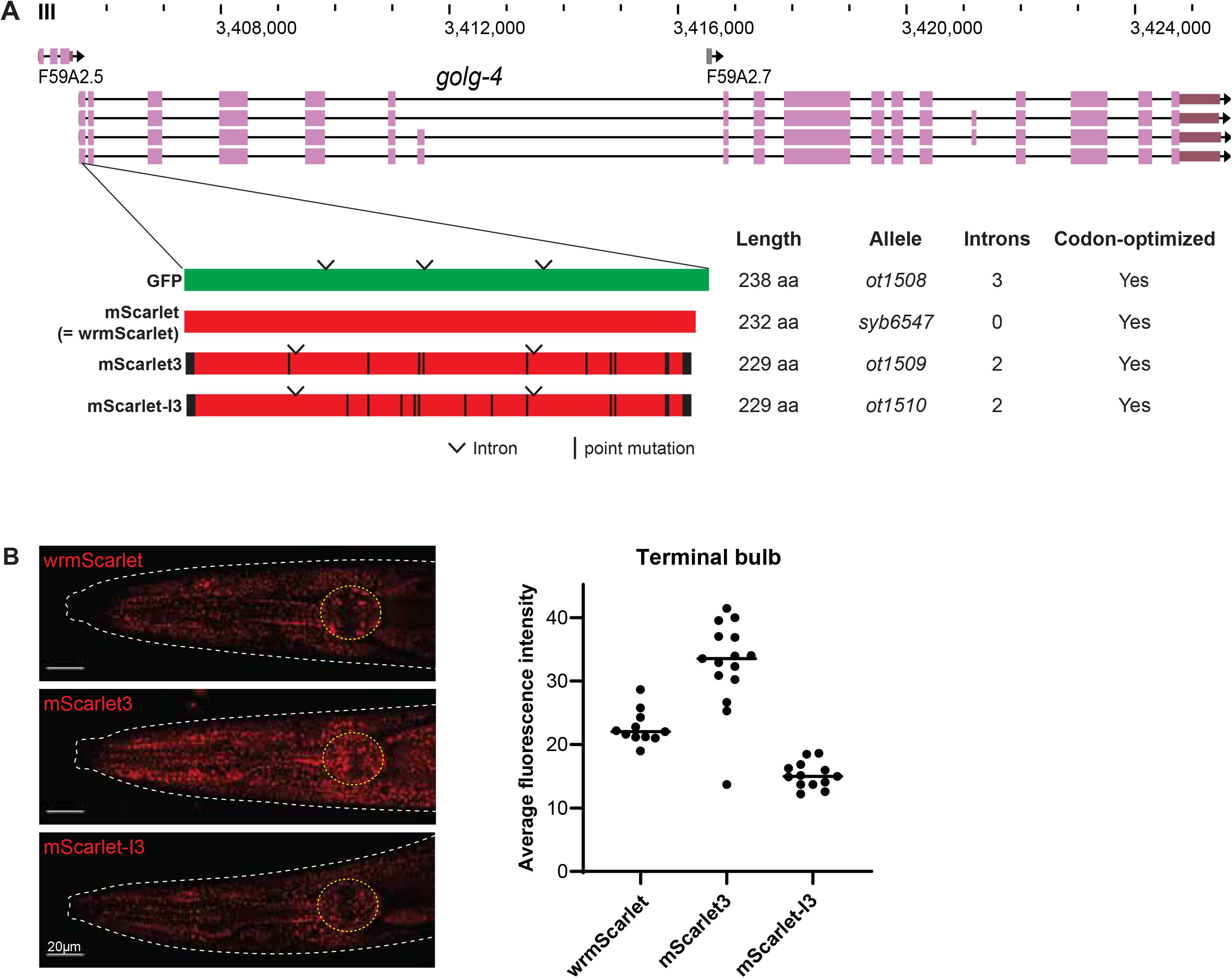
Analogous CRISPR-mediated insertion of fluorophores at the N-terminus of the GOLG-4 locus. **A:** WormBase genome browser snapshot of the *golg-4* locus and the site of CRIPSR fluorophore insert. Codon-optimized fluorophores are schematized below: derivatives of mScarlet – mScarlet3 and mScarlet-I3 – show amino acid mutations relative to mScarlet in black. GFP cloned from pPD95.75 expression plasmid is also included for comparison. Introns are also annotated and were added as recommended by the MPI *C. elegans* codon adapter (REDEMANN *et al*. 2011). **B:** Representative expression of red fluorophore-tagged GOLG-4 in the head is shown. Average fluorescence intensity of the pharyngeal terminal bulb of individual animals is quantified and plotted on the right, and the median values are indicated.

In all three reporter strains, tagged GOLG-4 is localized to puncta throughout the animal and is particularly bright in the gland cells of the posterior pharyngeal bulb (**Figure 1B**). To directly compare the brightness of these RFPs expressed *in vivo*, we quantified average fluorescence intensity of the posterior bulb in these CRISPR-tagged young adult worms. We find that mScarlet3 is about 50% brighter than wrmScarlet, while mScarlet-I3 is about 30% dimmer. These results correlate with *in vitro* characteristics of these fluorophores (GADELLA *et al*. 2023).

### mScarlet-I3 signal accumulates at comparable rates to GFP

We compared the maturation time of the RFPs against GFP by visualizing GOLG-4 accumulation in the developing *C. elegans* embryo. Embryos expressing fluorophore-tagged GOLG-4 were imaged at specific stages over the course embryo development from the 4-cell stage to the 4-fold stage, a period of about 9 hours, and compared to GOLG-4 similarly tagged with GFP, derived from a commonly used *C. elegans* expression plasmid from the Fire vector kit, pPD95.75 (**Figure 1A**).

We found that GFP::GOLG-4 puncta were detectable at low levels in the 4-cell embryo, and accumulates steadily at each subsequent stage captured throughout embryogenesis to the 4-fold stage (**Figure 2**). In contrast, wrmScarlet::GOLG-4 was not visible for several hours in the early embryo. wrmScarlet signal accumulates extremely slowly, is detected at low levels after onset of expression, and remains weakly expressed through the end of the time course. These observations are consistent with test data from mammalian cells, where mScarlet has a maturation time of nearly 3 hours at 37°C (GADELLA *et al*. 2023).

**Figure 2.**
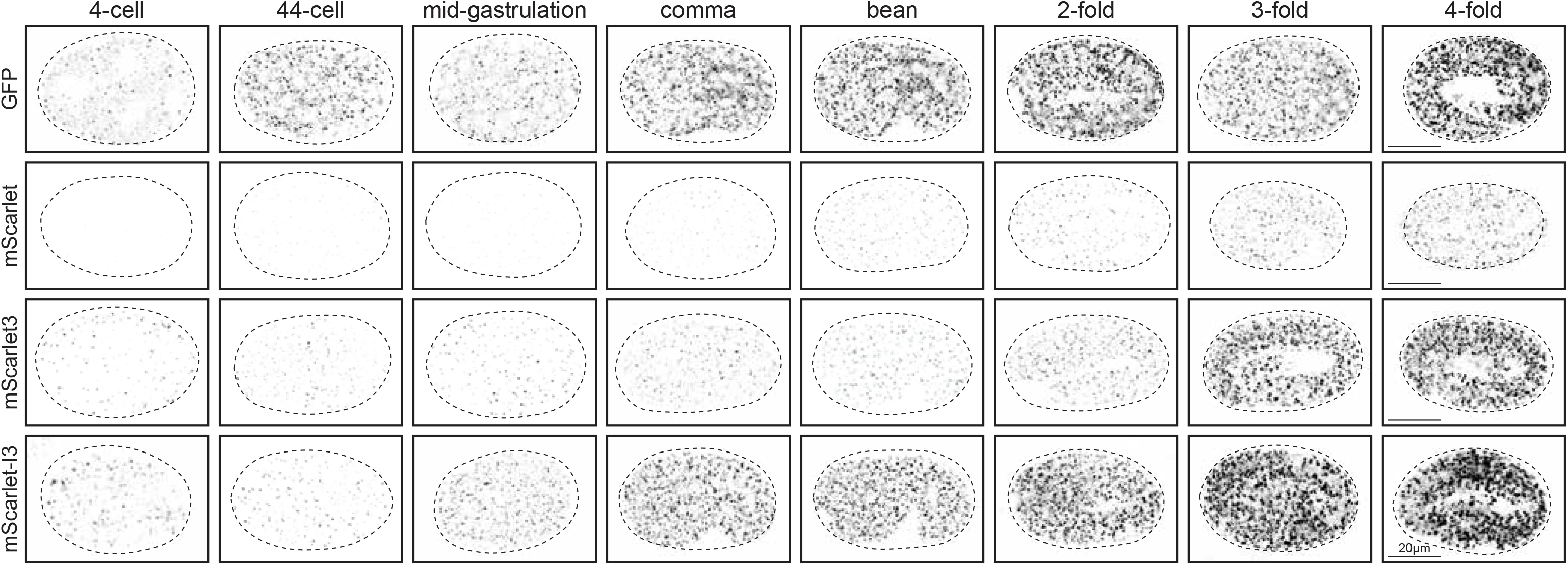
Expression of fluorophore-tagged GOLG-4 through embryonic development. Representative images of single focal plane through the middle of each embryo are shown, comparing the expression dynamics between the mScarlet variants and GFP. Strains are those shown in Figure 1. For comparison to GFP, imaging setting were chosen to match the fluorescence intensity of GFP::GOLG-4 to that of mScarlet-I3::GOLG-4 in the adult, quantified in Figure 1. Eight embryo developmental stages from the 4-cell stage to the 4-fold stage, spanning most the embryogenesis, are represented in columns and compared across fluorophores, in rows. Embryo stages were determined by DIC (not shown), and eggshells are traced in dashed lines.

Despite being a brighter fluorophore than wrmScarlet with a much faster maturation time in cell lines, mScarlet3::GOLG-4 fluorescence was also extremely slow to accumulate, detected only at low levels until the 3-fold stage (**Figure 2**). Evidently, the vast improvement of mScarlet3 maturation time – 37 minutes in mammalian cells (GADELLA *et al*. 2023) – had limited effects on expression dynamics in *C. elegans*.

In stark contrast, the onset of mScarlet-I3::GOLG-4 expression is comparable to GFP::GOLG-4. Like the GFP reporter allele, it is also visible at the 4-cell stage, and also accumulates readily and steadily over the captured time points through embryo development (**Figure 2**). Moreover, mScarlet-I3 signal intensity surpasses that of wrmScarlet and mScarlet3 by the 3-fold stage, despite being a dimmer fluorophore. Indeed, mScarlet-I3::GOLG-4 signal accumulates to higher levels much faster than its RFP counterparts. We conclude that in *C. elegans* mScarlet-I3 is a bright, rapidly-maturing RFP that is able to capture protein expression dynamics with minimal delay, compared to GFP.

### mScarlet-I3 behavior in other cells and subcellular locales

To independently validate these findings in different cellular and subcellular contexts, we tagged another locus, *pks-1*, encoding a polyketide synthase (FENG *et al*. 2021), with GFP or mScaret-I3. In this case, we did not fuse the fluorophores directly to the gene, but used a standard polycistronic cassette approach in which the fluorophores were separated from the tagged gene with a trans-splicing SL2 signal and targeted to the nucleus by addition of a histone 2B (*his-44*) cassette (**Figure 3A**). Late larval stage animals carrying *pks-1::SL2::mScarlet-I3::H2B* or *pks-1::SL2::GFP::H2B* show fluorophore expression in the nuclei of the CAN neuron pair (consistent with previous findings; (FENG *et al*. 2021)) and in cells of the alimentary system, including intestinal cells as well as pharyngeal cells (**Figure 3B-C**). Notably, pharyngeal cells showed brighter signal with the mScarlet-I3 reporter compared to the GFP reporter; the mScarlet-I3 signal also appeared crisper because of lesser background fluorescence (**Figure 3B**). In the intestine, the signal-to-noise ratio is even more improved with the mScarlet-I3 reporter since the characteristic autofluorescent puncta of the intestine in the green channel are nearly absent when imaging in the red channel. As a result, the adjacent CAN nuclei in the midbody of the worm are much more easily discernable when marked with mScarlet-I3 (**Figure 3B**).

**Figure 3.**
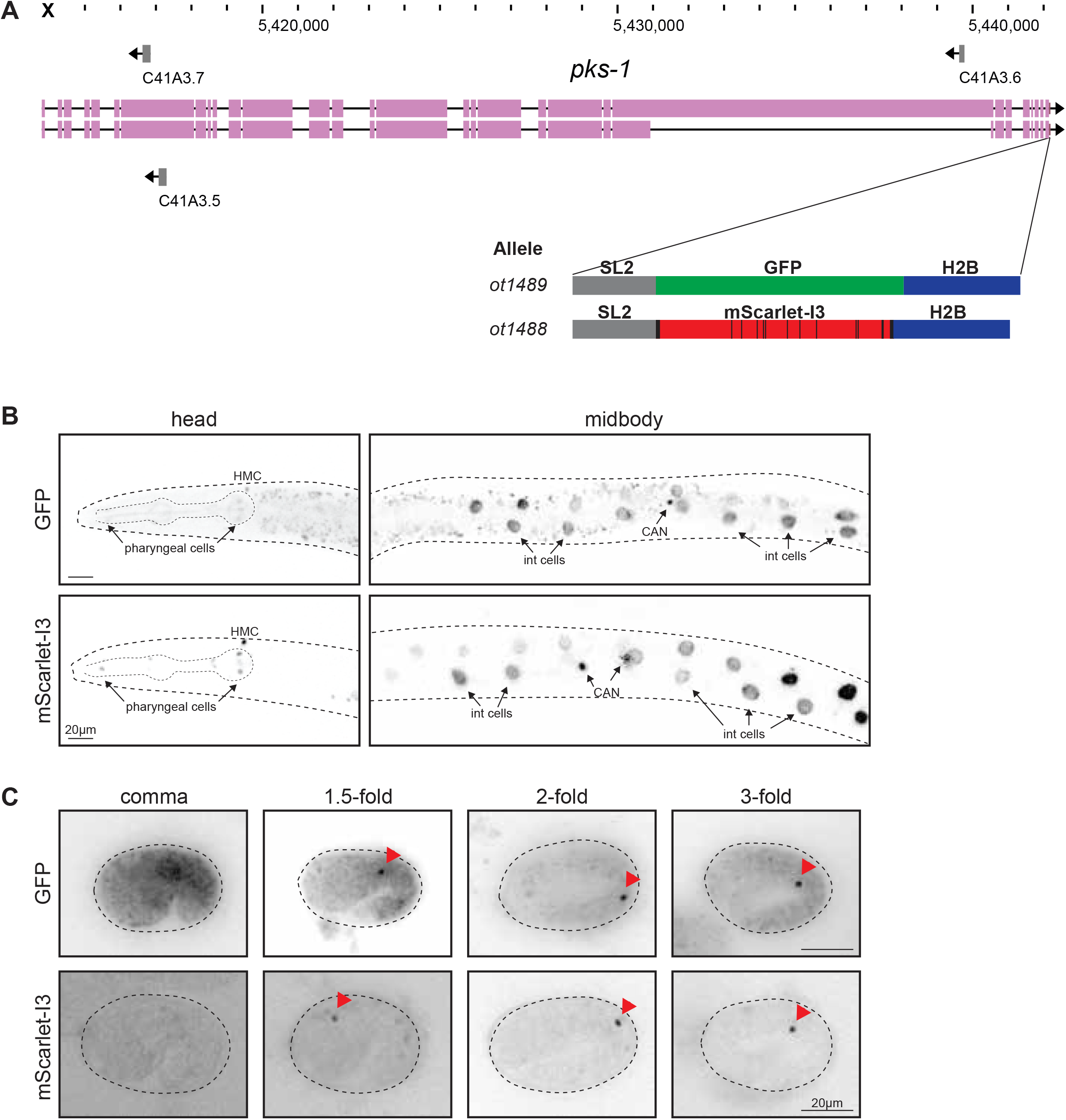
Expression of *pks-1::SL2::GFP::H2B* and *pks-1::SL2::mScarlet-I3::H2B* during embryo development. **A:** WormBase genome browser snapshot of the *psk-1* locus and the site of CRISPR-mediated insertion at the C-terminus, after the stop codon. Schematic of fluorophore constructs are shown below. **B:** Expression of fluorophores in the L3 larval head and midbody. Both GFP and mScarlet-I3 reporters are expressed in the nuclei of CANL/R and intestinal (int) cells. Lower expression is also observed in several nuclei within the pharynx. An additional nucleus in the head, presumed to be the head mesodermal cell (HMC), is also labeled. Images are of a single plane containing one of the two CAN nuclei in focus. **C:** Expression of fluorophores in various stages of the embryo. Focal plane containing one of the two nuclei is shown for each embryo. Fluorescent signal can be visualized from the comma stage onwards, for both GFP and mScarlet-I3 constructs (red arrowheads). Note the *pks-1* reporter constructs specifically label CAN nuclei at these embryonic stages shown.

Next, these animals were analyzed for onset of fluorophore expression in the embryo and found to display comparable expression dynamics: fluorescently-labeled CAN nuclei are not visible at the bean stage with either the GFP or mScarlet-I3 fluorophore, but become clearly detectable shortly after by the comma stage and through the rest of embryo development (**Figure 3C)**. We again observe much less background autofluorescence when imaging the mScarlet-I3 reporter compared to the GFP reporter, especially at the earlier embryonic stages. (**Figure 3C**).

Taken together, mScarlet-I3 appears superior to GFP in terms of increased signal intensity, decreased background particularly in the intestines, while exhibiting a comparable maturation time to GFP.

### mScarlet-I3::UNC-75 as a red pan-neuronal marker

As an additional case study, we translationally tagged *unc-75*, a CELF family RNA-binding protein expressed specifically in all neuronal nuclei but no other cell types (LORIA *et al*. 2003), with both GFP and mScarlet-I3 (**Figure 4A**) and characterized onset of expression of the resulting reporter alleles. We observe pan-neuronal nuclear expression of both fluorophores throughout the adult worm. Consistent with the above observations, we also observe improved signal-to-noise ratio of mScarlet-I3 reporter compared to the GFP reporter, particularly resulting from decreased autofluorescence in the intestines associated with a longer excitation laser when imaging mScarlet-I3 (**Figure 4B**).

**Figure 4.**
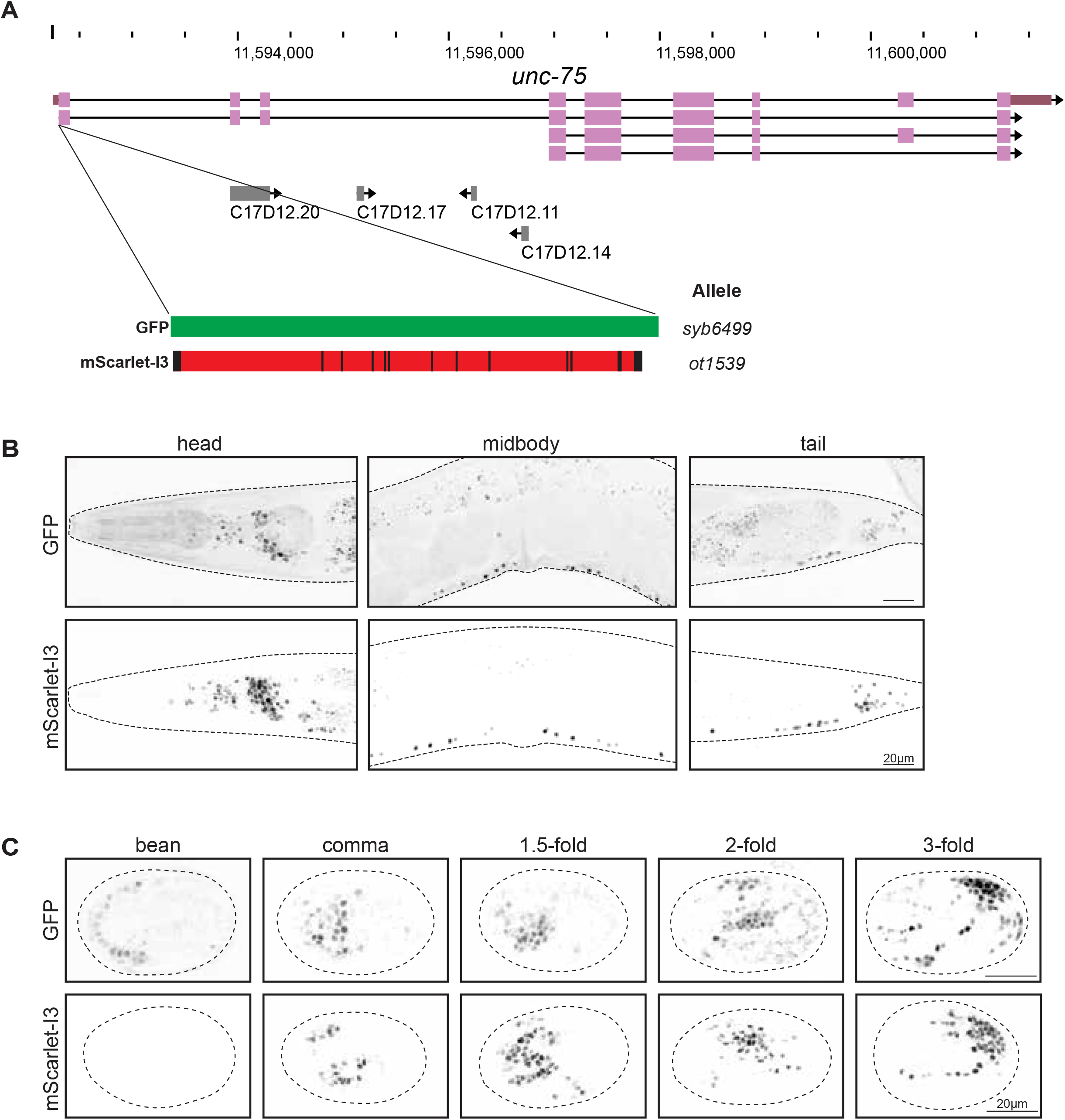
Expression of fluorophore-tagged UNC-75, comparing GFP to mScarlet-I3. **A:** WormBase genome browser snapshot of the *unc-75* locus, showing the site of CRISPR-mediated insertion at the N-terminus. Fluorophore constructs are the same as in previous figures. The fluorophores were inserted at the N-terminus to avoid disruption of a C-terminal nuclear localization sequence (LORIA *et al*. 2003). **B:** Expression of fluorophore-tagged UNC-75 in the adult worm. Images are maximum intensity projections through the head, midbody and tail sections, capturing dense regions of neuron nuclei. **C:** Expression of fluorophore-tagged UNC-75 at five stages in embryo development, determined by DIC. Single Z planes through the approximate midpoint of each embryo are shown, and eggshells are traced with dashed lines.

We observe the earliest onset of GFP::UNC-75 expression at low levels around the bean stage (**Figure 4C**). GFP signal accumulates quickly in subsequent embryonic stages. In comparison, mScarlet-I3::UNC-75 levels were just below detection threshold at the bean stage, but become visible shortly after, accumulating to comparable levels by the comma stage (**Figure 4C**). Hence, in this case, expression of mScarlet-I3 slightly lags behind that of GFP. However, we also note that like in adults, mScarlet-I3::UNC-75 expression in the embryo has less non-specific background fluorescence signal and better signal-to-noise ratio from the onset of visible expression compared to GFP::UNC-75.

## Conclusions

We described here that mScarlet-I3 is the fastest maturing mScarlet-based fluorescent marker currently known in *C. elegans*, making it the prime choice for usage if maturation time is an important factor. mScarlet-I3 is not quite as bright as mScarlet3, but given its overall brightness combined with low background fluorescence in the red channel, this only becomes an issue if gene or protein expression are particularly low. mScarlet3 may be more useful for visualizing especially lowly expressed genes when expression dynamics are not of concern.

## ACKNOWLEDGEMENTS

We thank Chi Chen for nematode injections and members of the Hobert lab for discussions and comments on the manuscript. This work was funded by NIH R01NS039996. Wen Xi Cao is supported by a Canadian Institutes of Health Research (CIHR) fellowship MFE-176611.

## DATA AVAILABILITY

All strains are being deposited at the CGC.

## AUTHOR CONTRIBUTIONS

WXC: Conceptualization, Data curation, Formal analysis, Investigation, Visualization, Writing – original draft preparation, DM: Conceptualization, Investigation, KP, MC: Investigation; OH: Conceptualization, Funding acquisition, Project administration, Supervision, Writing – review and editing.

## Supplemental file S1: Fluorophore sequences

The *C. elegans* codon-optimized mScarlet-3 and mScarlet-I3 sequences below have been cloned into the pPD95.75 backbone in place of GFP and will be deposited to Addgene with the following plasmid names:

pWXC077: mScarlet3

pWXC078: mSarlet-I3

Overhang sequences used to replace GFP in pPD95.75 were:

~~~
5’-gacccttggagggtaccggtagaaaaa-3’
5’-cattcgtagaattccaactg-3’
~~~

**XXXX** `coding sequence`

~~~
xxxx introns
~~~

**>mScarlet3 ATGGACTCCACCGAGGCCGTCATCAAGGAGTTCATGCGTTTCAAGGTCCACATGGAGGGATCCA TGAACGGACACGAGTTCGAGATCGAGGGAGAGGGAGAGGGACGTCCATACGAGGGAACCCAAAC CGCCAAGCTCCGTGTCACCAAG**gtaagtttaaacatatatatactaactaaccctgattattta aattttcag**GGAGGACCACTCCCATTCTCCTGGGACATCCTCTCCCCACAATTCATGTACGGAT CCCGTGCCTTCACCAAGCACCCAGCCGACATCCCAGACTACTGGAAGCAATCCTTCCCAGAGGG ATTCAAGTGGGAGCGTGTCATGAACTTCGAGGACGGAGGAGCCGTCTCCGTCGCCCAAGACACC TCCCTCGAGGACGGAACCCTCATCTACAAGGTCAAGCTCCGTGGAACCAACTTCCCACCAGACG GACCAGTCATGCAAAAGAAGACCATGGGATGGGAGGCCTCCACCGAGCGTCTCTACCCAGAGGA CGTCGTCCTCAAG**gtaagtttaaacagttcggtactaactaaccatacatatttaaattttcag **GGAGACATCAAGATGGCCCTCCGTCTCAAGGACGGAGGACGTTACCTCGCCGACTTCAAGACCA CCTACCGTGCCAAGAAGCCAGTCCAAATGCCAGGAGCCTTCAACATCGACCGTAAGCTCGACAT CACCTCCCACAACGAGGACTACACCGTCGTCGAGCAATACGAGCGTTCCGTCGCCCGTCACTCC ACCGGAGGATCCGGAGGATCCTAA**

**>mScarlet-I3 ATGGACTCCACCGAGGCCGTCATCAAGGAGTTCATGCGTTTCAAGGTCCACATGGAGGGATCCA TGAACGGACACGAGTTCGAGATCGAGGGAGAGGGAGAGGGACGTCCATACGAGGGAACCCAAAC CGCCAAGCTCAAGGTCACCAAG**gtaagtttaaacatatatatactaactaaccctgattatttaaattttcag**GGAGGACCACTCCCATTCTCCTGGGACATCCTCTCCCCACAATTCATGTACGGAT CCCGTGCCTTCATCAAGCACCCAGCCGACATCCCAGACTACTGGAAGCAATCCTTCCCAGAGGG ATTCAAGTGGGAGCGTGTCATGATCTTCGAGGACGGAGGAACCGTCTCCGTCACCCAAGACACC TCCCTCGAGGACGGAACCCTCATCTACAAGGTCAAGCTCCGTGGAGGAAACTTCCCACCAGACG GACCAGTCATGCAAAAGCGTACCATGGGATGGGAGGCCTCCACCGAGCGTCTCTACCCAGAGGA CGTCGTCCTCAAG**gtaagtttaaacagttcggtactaactaaccatacatatttaaattttcag **GGAGACATCAAGATGGCCCTCCGTCTCAAGGACGGAGGACGTTACCTCGCCGACTTCAAGACCACCTACAAGGCCAAGAAGCCAGTCCAAATGCCAGGAGCCTTCAACATCGACCGTAAGCTCGACATCACCTCCCACAACGAGGACTACACCGTCGTCGAGCAATACGAGCGTTCCGTCGCCCGTCACTCCACCGGAGGATCCGGAGGATCCTAA**

**>wrmScarlet GTCAGCAAGGGAGAGGCAGTTATCAAGGAGTTCATGCGTTTCAAGGTCCACATGGAGGGATCCATGAACGGACACGAGTTCGAGATCGAGGGAGAGGGAGAGGGACGTCCATACGAGGGAACCCAAACCGCCAAGCTCAAGGTCACCAAGGGAGGACCACTCCCATTCTCCTGGGACATCCTCTCCCCACAATTCATGTACGGATCCCGTGCCTTCACCAAGCACCCAGCCGACATCCCAGACTACTACAAGCAATCCTTCCCAGAGGGATTCAAGTGGGAGCGTGTCATGAACTTCGAGGACGGAGGAGCCGTCACCGTCACCCAAGACACCTCCCTCGAGGACGGAACCCTCATCTACAAGGTCAAGCTCCGTGGAACCAACTTCCCACCAGACGGACCAGTCATGCAAAAGAAGACCATGGGATGGGAGGCCTCCACCGAGCGTCTCTACCCAGAGGACGGAGTCCTCAAGGGAGACATCAAGATGGCCCTCCGTCTCAAGGACGGAGGACGTTACCTCGCCGACTTCAAGACCACCTACAAGGCCAAGAAGCCAGTCCAAATGCCAGGAGCCTACAACGTCGACCGTAAGCTCGACATCACCTCCCACAACGAGGACTACACCGTCGTCGAGCAATACGAGCGTTCCGAGGGACGTCACTCCACCGGAGGAATGGACGAGCTCTACAAG**

~~~
(no start or stop included because this sequence was inserted at the C-terminus, before the stop codon, of the golg-4 gene)
~~~

**>GFP** `(from pPD95.75)`

**ATGAGTAAAGGAGAAGAACTTTTCACTGGAGTTGTCCCAATTCTTGTTGAATTAGATGGTGATG TTAATGGGCACAAATTTTCTGTCAGTGGAGAGGGTGAAGGTGATGCAACATACGGAAAACTTAC CCTTAAATTTATTTGCACTACTGGAAAACTACCTGTTCCATGG**gtaagtttaaacatatatata ctaactaaccctgattatttaaattttcag**CCAACACTTGTCACTACTTTCTGTTATGGTGTTC AATGCTTCTCGAGATACCCAGATCATATGAAACGGCATGACTTTTTCAAGAGTGCCATGCCCGA AGGTTATGTACAGGAAAGAACTATATTTTTCAAAGATGACGGGAACTACAAGACACGT**aagttt aaacagttcggtactaactaaccatacatatttaaattttcaggt**GCTGAAGTCAAGTTTGAAG GTGATACCCTTGTTAATAGAATCGAGTTAAAAGGTATTGATTTTAAAGAAGATGGAAACATTCT TGGACACAAATTGGAATACAACTATAACTCACACAATGTATACATCATGGCAGACAAACAAAAG AATGGAATCAAAGTT**gtaagtttaaacatgattttactaactaactaatctgatttaaattttc ag**AACTTCAAAATTAGACACAACATTGAAGATGGAAGCGTTCAACTAGCAGACCATTATCAACA AAATACTCCAATTGGCGATGGCCCTGTCCTTTTACCAGACAACCATTACCTGTCCACACAATCT GCCCTTTCGAAAGATCCCAACGAAAAGAGAGACCACATGGTCCTTCTTGAGTTTGTAACAGCTG CTGGGATTACACATGGCATGGATGAACTATACAAATAG**

